# Sub-lethal insecticide exposure affects host biting efficiency of *Kdr*-resistant *Anopheles gambiae*

**DOI:** 10.1101/653980

**Authors:** Malal M Diop, Fabrice Chandre, Marie Rossignol, Angélique Porciani, Mathieu Chateau, Nicolas Moiroux, Cédric Pennetier

**Affiliations:** MIVEMEC, Univ Montpellier, CNRS, IRD, Montpellier, France; Institut de Recherche en Sciences de la Santé (IRSS), Bobo-Dioulasso, Burkina Faso; Institut Pierre Richet (IPR), Bouaké, Côte d’Ivoire

**Keywords:** pyrethroid insecticides, long-lasting treated nets, malaria, probing, blood meal, prediuresis, resistance, kdr

## Abstract

The massive use of insecticide-treated nets (ITNs) has drastically changed the environment for malaria vector mosquitoes, challenging their host-seeking behaviour and biting success. Here, we investigated the effect of a brief exposure to an ITN on the biting behaviour of *Anopheles mambiae* mosquitoes and the interaction between such behaviour and the *kdr* mutation that confers resistance to pyrethroids. To this aim, we developed a video assay to study the biting behaviour of mosquitoes with similar genetic background, but different *kdr* locus genotypes (SS i.e. homozygous susceptible, RS i.e. heterozygous and RR i.e. homozygous resistant), after a brief exposure to either control untreated nets or one of two types of pyrethroid-treated nets (deltamethrin or permethrin). In presence of untreated nets, the *kdr* mutation did not influence mosquito blood feeding success but caused differences in feeding and prediuresis durations and blood meal size. Exposure to deltamethrin ITN decreased the blood feeding success rate of RR and RS mosquitoes, whereas in presence of permethrin ITN, the *kdr* mutation increased the blood-feeding success of mosquitoes. Exposure to the two types of pyrethroid-treated nets reduced feeding duration, prediuresis duration and blood meal size of all three genotypes. Our study demonstrates a complex interaction between insecticide exposure and the *kdr* mutation on the biting behavior of mosquitoes, which may substantially impact malaria vector fitness and disease transmission.

## Introduction

Malaria vector mosquitoes become infected by and then can transmit *Plasmodium spp*. parasites during blood meals. *Plasmodium spp*. transmission is strongly dependent on the mosquito biting rate on humans, as formalized by the Ross-MacDonald model of malaria (1). The behaviour and host preferences of blood-feeding mosquitoes are influenced by several factors, primarily their genetic background (2–5), but also environmental factors, such as host diversity and availability (6,7), and the presence/absence of physical and chemical barriers, such as Insecticide-Treated Nets (ITNs) (8–10).

ITNs should hamper the contacts between humans and nocturnal, anthropophilic (i.e. that bite humans preferentially) and endophagic mosquitoes (i.e. that bite preferentially indoors), such as *An. mambiae* (11). The widespread use of pyrethroid (PYR) insecticides in public health (i.e. ITNs or indoor residual sprayings), crop protection and other selective pressures (such as pollutants in mosquito environment) drove adaptations in malaria vectors to reduce the insecticidal effect (12–17).

Physiological PYR resistance involves two main mechanisms: (i) metabolic resistance, due to quantitative or qualitative changes in detoxification enzymes (cytochrome P450 monooxygenases, esterases and glutathione S-transferases), and (ii) target site resistance, due to non-synonymous mutations in the voltage-gated sodium channels that are called knock-down resistance (*kdr*) mutations (12,18).

Mosquitoes can reduce vector control tools efficacy also through behavioural adaptations. In areas of sub-Saharan Africa where large-scale vector control programmes have been implemented, several observations suggest that malaria vectors can avoid contacts with ITNs or insecticides on walls by modulating their host-feeding activity. For instance, following the large scale distribution of ITNs in Benin, some An. funestus populations started to feed predominantly at dawn/early in the morning or in broad daylight (19,20), when most people are outside the nets (20). Behavioural modulation might also be influenced by physiological resistance mechanisms. Indeed, experimental hut trials showed that in *An. mambiae*, indirect behavioural indicators in response to ITN presence (deterrence and induced exophily, i.e. the proportion of mosquitoes that exit early and are found in exit traps relative to the untreated hut) are related to the physiological tolerance to insecticide (21). Moreover, a study demonstrated that *kdr* homozygous resistant mosquitoes have longer contacts with ITNs than homozygous susceptible mosquitoes, which are more excited by PYR irritant effect (22). Altogether, these findings indicate that insecticide treatments could affect the behaviour of malaria vectors. However, the effects of insecticide exposure and *kdr* mutations on the biting activity of *An. mambiae* remain poorly investigated.

The last step of the mosquito host-seeking behaviour after reaching a host protected by an ITN is biting for taking a blood meal. During the host-seeking phase and the penetration through a hole in the net, mosquitoes can be exposed to sub-lethal doses of insecticide (23–25). Such doses do not cause death, but can have several physiological or behavioural effects on host-seeking mosquitoes (26). For instance, sub-lethal doses of insecticide can affect the feeding and reproductive behaviour of some blood-sucking insects (27). As the PYR insecticide on ITNs acts on specific sites in the mosquito nervous system, it might alter some physiological processes involved in the biting behaviour of malaria vectors. There is currently scarce literature on this subject (28).

In the present study, we investigated whether pre-exposure to an ITN modulates the mosquito ability to take a blood meal by using experimental conditions that mirror the exposure to insecticide occurring when a mosquito passes through an ITN after having located a host. We also assessed whether the *kdr* mutation (L1014F) modifies blood feeding success and biting behaviour of *An. mambiae*.

## Methods

### Ethical Considerations

This study was carried out in strict accordance with the recommendations of Animal Care and Use Committee (29) named “Comité d’éthique pour l’expérimentation animale; Languedoc Roussillon” and the protocol was approved by this committee (CEEA-LR-13002 for the rabbits).

### Mosquito strains and rearing procedures

Two mosquito laboratory strains were used for this study. One is the insecticide-susceptible Kisumu strain (KISUMU1, MRA-762, VectorBase stable ID VBS0000026 on vectorbase.org), isolated in Kenya in 1975. This strain is PYR-susceptible and homozygous (SS) for the L1014 codon. The second one is the Kdr-kis strain that is PYR-resistant and homozygous (RR) for the L1014F *kdr* mutation. The Kdr-kis strain was obtained by introgression of the *kdr* allele (L1014F) into the Kisumu genome through 19 successive back-crosses between Kisumu and VKPer (30). The VKPer strain, originated from a rice growing area named Vallée du Kou, less than 40 kms north of Bobo-Dioulasso (Burkina Faso) was used to obtain Kdr-kis. VKPer displayed the same expression level of metabolic resistance enzyme as Kisumu (31). Both Kisumu and Kdr-kis strains are maintained at the insectary of IRD (Institut de Recherche pour le Développement) – WHO collaborating center FRA-72 in Montpellier, France.

Polymorphisms between Kisumu and Kdr-kis strains are expected to be restricted in the flanking region of the *kdr* allele (15 cM for 19 backcrossing generations (32)) and the observed phenotypes are therefore expected to be associated to this genetic area.

Heterozygous individuals (RS) for the L1014F *kdr* mutation were obtained by crossing once Kisumu SS females (F1 progeny) with Kdr-kis RR males. Therefore, the three genotypes have a common genetic background for most of their genome.

Mosquitoes were reared at 27 ± 1°C, 70-80% relative humidity under a 12h:12h (light : dark) photoperiod in the insectary and fed on rabbits. Mravid females were allowed to lay eggs on wet filter paper inside mesh-covered cages. Eggs were dispensed in plastic trays containing osmotic water. Larvae were kept in trays and fed with TetraMin® fish food. Pupae were removed and allowed to emerge inside 30×30×30cm cages. After emergence, adults could feed ad libitum on 10% sucrose solution.

On each experimental day, an average of 8 (2 to 16 according to the insectary production) female mosquitoes (7 to 9 days old) never fed with blood were randomly collected from the rearing cages and placed in cups covered by gauze. By using 7-9 days old *An. mambiae*, we expected to enhance the probability that these females were inseminated (33) and physiologically active for host-seeking (34). However, mated status was not checked.

Mosquitoes were starved the day before the experiment because sucrose inhibits blood avidity in mosquitoes (35).

### Insecticide exposure

To simulate the contact with an ITN that may occur before the vector finds a way to reach a host, starved females were individually exposed to PYR insecticide-treated netting using an experimental setup in which insects can only walk or stand on the net surface and that was initially designed to measure the median knock down time (36). We used three net types (i.e. treatments): an untreated net (negative control, hereafter UTN), an Olyset Net® (impregnated with 1000 mg/m2 of permethrin, hereafter Olyset) and a PermaNet® 2.0 (coated with 55 mg/m2 of deltamethrin, hereafter PermaNet). Based on previous experiments (25), each mosquito was exposed for 30 seconds. This is the median time of contact with permethrin-treated netting before PYR-susceptible anopheles (SS genotype) locate a hole to reach the host (25). After exposure, a 1-min latency period was observed before releasing the insect in the behavioural assay setup.

### Behavioural assay

Experiments were conducted in a biting behavioural assay setup designed for video recording the biting behaviour of the tested mosquitoes (Figure 1). The setup is made of a foam board and composed of an observation tunnel (OT) and an observation zone (OZ) separated by a transparent plastic (TP). The OZ is a triangular prism, closed on its base by a removable paper sheet with a 1-cm diameter hole. The hole allows mosquitoes to bite the ear of a rabbit (R) that is maintained immobile in a restraining cage (RC) to limit ear movement during blood feeding. The same rabbit was used during all experiments. The experimental room was faintly illuminated with a compact fluorescent lamp bulb placed at 15 cm from the OZ to allow the acquisition of the biting behaviour sequence by the digital video system.

**Figure 1.**
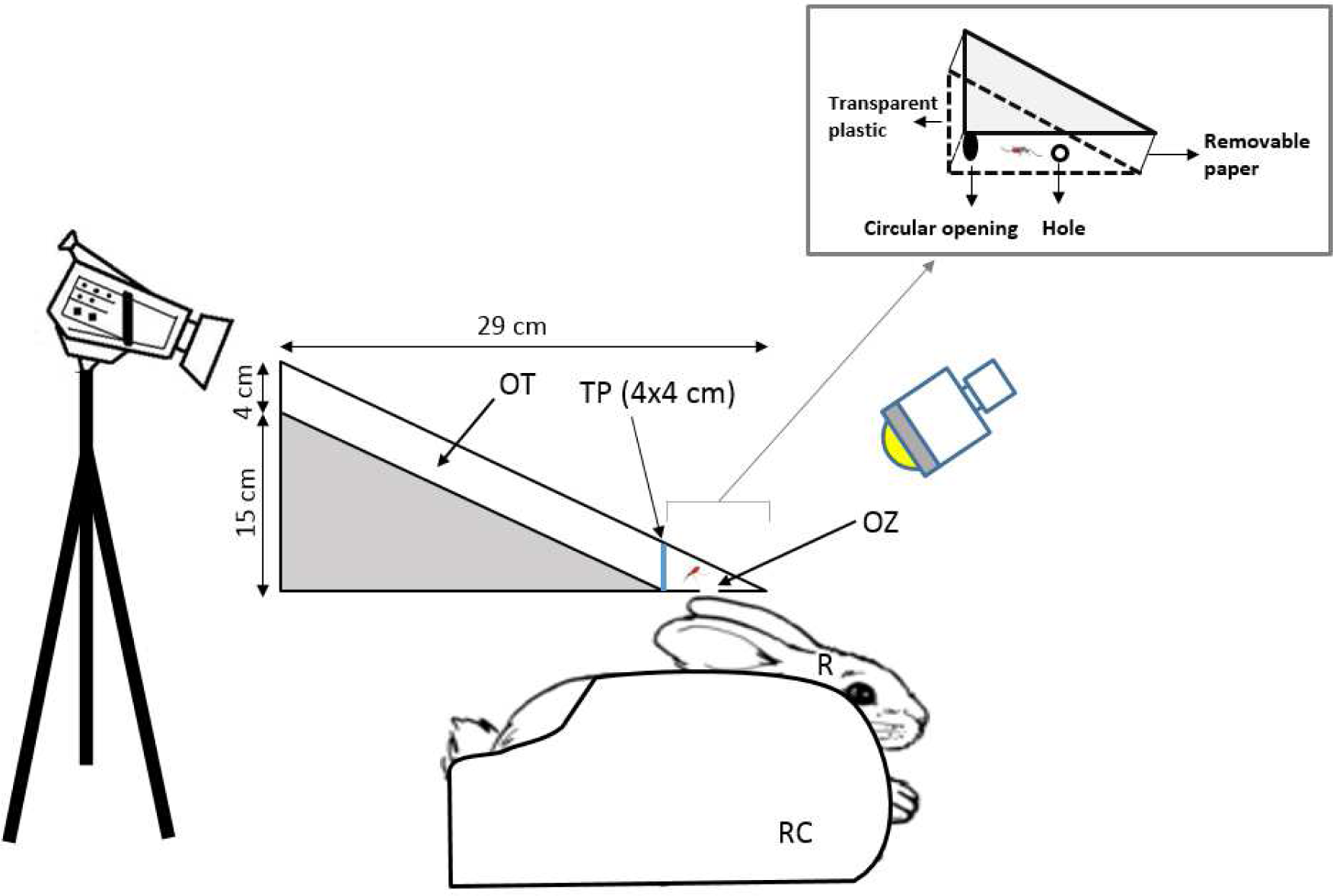
Experimental set-up to monitor the biting behavioural sequence. OT: Observational tunnel, TP: Transparent plastic, OZ: Observational zone (mosquito biting the rabbit ear), R: Rabbit, RC: Restraining cage

For each trial, a mosquito was individually released inside the behavioural assay through a circular opening (CO) located on its lateral face. Cotton was used to plug the CO after releasing. The number of released individual anopheles per genotype and treatment ranged from 43 to 86. Each mosquito was filmed for 10 minutes using a Sony® Digital HD Video Camera (HDR-XR550) placed on the top of the OT (Figure 1). The video camera was connected to a computer screen placed outside the experimental room to allow real-time monitoring of each mosquito, to avoid any disturbance of the rabbit and interference of the experimenter on the mosquito behaviour. The MPEM-2 recordings (PAL video: 720×576 pixels at 25 frames/s) were analysed using the behavioural event recording program EVENT01, version 1.2.4 (©R.D. Collins and M.K. Tourtellot, 1993-2002).

The experimenter used latex gloves to avoid any contamination by human skin odours. The OZ was cleaned with ethanol and the removable paper was changed after each mosquito was tested to avoid any contamination by insecticide residues between behavioural assays. The feeding experiments were done under insectary conditions (temperature at 27 ± 1°C, and relative humidity 70-80%), in a dark room during the night (according to the Light:Dark photoperiod of the insectary).

### Feeding success and behavioural parameters

As exposure to insecticide can induce a knockdown (KD) effect during the trial, mosquitoes were recorded as KD, if they were lying on their side or their back. A mosquito (whatever its KD phenotype) was scored as successful if it was fed at the end of the 10-min trial (whatever the amount of blood it took) and unsuccessful if it did not. After the trial, successfully fed mosquitoes were stored at −35°C for measuring the blood meal size (see below).

Analysis of the acquired images allowed quantifying the following variables in fed mosquitoes: (i) number of probing events, (ii) probing duration (the time from the introduction of the stylet fascicule into the rabbit skin to blood appearance in the mosquito abdomen), (iii) feeding duration (from the blood appearance in the abdomen to the beginning of the stylet fascicule withdrawal), and (iv) prediuresis duration (the time during which excretion of rectal fluid (plasma, water, metabolic wastes) is observed as red bright drops during feeding and after proboscis withdrawal).

### Blood meal volume measurement

The blood intake was evaluated by quantifying the haemoglobin amount, as described by Briegel *et al*. (37). Each engorged mosquito was stored in one 1.5 ml Eppendorf tube at −35°C. Then, the whole abdomen was ground in the presence of 0.5 ml of Drabkin’s reagent until it was completely disintegrated. Haemoglobin then reacted with the Drabkin’s reagent and was converted into haemoglobin cyanide (HiCN). Samples were incubated at room temperature (25°C) for 20 min and then a chloroform solution was added in each tube. Samples were centrifuged at 5600 rpm for 5 min and the aqueous supernatant (containing HiCN) was placed in a new 1.5 ml Eppendorf tube. The absorbance was read at a wavelength of 550 nm using a microplate spectrophotometer. Two replicates of the reading were done for each mosquito and their absorbance values were averaged. A sample of the rabbit blood was used as control for calibration curves.

As the blood meal volume is correlated with the mosquito size (38), the blood meal volume of all mosquitoes from the same batch was divided by the average weight of five randomly selected mosquitoes from the same rearing cage. The resulting ratio was expressed in µL of blood per µg of weight and called weighted blood meal volume or blood meal size.

## Statistical analysis

All statistical analyses were performed using the R software, version 3.5 (39).

We analysed the feeding success (coded as 1 for fed mosquitoes and 0 for unfed ones) with a binomial logistic mixed-effect model using function ‘glmmTMB’ in the ‘glmmTMB’ package (40). We analysed the number of probing events with a zero-truncated negative binomial mixed-effect model using function ‘glmmTMB’. We analysed durations (probing, feeding and prediuresis) with a mixed effect Cox proportional hazard model using function ‘coxme’ of the ‘coxme’ package (41). We analysed weighted blood meal size with a linear mixed effect model using function ‘lmer’ in the ‘lme4’ package (42). All models included the *kdr* genotypes (SS, RS or RR), type of pre-exposure (UTN, Permethrin or Deltamethrin) and their interactions as fixed terms explanatory variables and the date of the experiment as a random intercept.

In order to discriminate between the effect of KD phenotype and other pleiotropic effects of the *kdr* mutation on feeding success and behavioural parameters, we fitted all previously described models on the dataset but for KD mosquitoes. To complement the later analysis, we tested the effect of the KD phenotype on feeding success. For this task, we added KD phenotype and its interactions as explanatory variables in the binomial model of feeding success fitted only on mosquitoes exposed to the insecticide treatments (Permethrin and Deltamethrin). This allowed us to compare feeding success between KD and non-KD mosquitoes of each genotype and treatment.

In order to assess the relationship between KD phenotype and the genotypes for the *kdr* mutation, KD (coded as 1 for KD mosquitoes and 0 for others) was analysed using a binomial model with the *kdr* genotypes (SS, RS or RR), type of exposure (Permethrin or Deltamethrin) and their interactions as fixed terms. Because the dataset showed data separation, we fitted the model using the bias-reduction method developed by Firth (43). We used the ‘brglmFit’ function of the ‘brglm2’ package (44) for this task.

We used Tukey’s post-hoc test to perform multiple comparisons among genotypes and treatments using ‘emmeans’ function (45). We computed Odds Ratios (OR) for the binomial models, Hazard Ratios (HR) for Cox models, Rate Ratios (RR) for the negative binomial models, Mean Differences (MD) for the linear models and their 95% confidence intervals.

We calculated the binomial confidence interval of feeding rates and knock-down rates with the Wilson’s score method using the ‘binconf’ command from the ‘Hmisc’ package (46).

Effect of feeding duration on blood meal size and prediuresis duration was analysed using a linear mixed effect model and a mixed effect cox proportional hazard model, respectively. Fixed terms included genotypes and treatment and interactions and date was set as a random intercept. We used function ‘emtrends’ of the ‘emmeans’ package (45) to obtain estimates and 95% confidence intervals of the marginal slopes of the trends (between feeding duration and blood meal size or prediuresis duration) for each genotypes and treatments.

Data and codes used for analyses and figures are available (47).

## Results

In total, 511 *An. mambiae* females (182 SS, 156 RS and 173 RR) were released individually for the study. The number of mosquitoes released, fed, unfed and knock-downed among genotypes and treatments are shown in Table 1.

**Table 1.** Numbers of mosquitoes released per genotype and treatment.

| Genotype | Treatment | N tested | Fed |  | Unfed |  |
| --- | --- | --- | --- | --- | --- | --- |
|  |  |  | KD | Non-KD | KD | Non-KD |
| SS | Untreated | 86 | 0 | 41 | 0 | 45 |
| SS | Permethrin | 50 | 5 | 5 | 33 | 7 |
| SS | Deltamethrin | 46 | 1 | 15 | 18 | 12 |
| RS | Untreated | 70 | 0 | 40 | 0 | 30 |
| RS | Permethrin | 43 | 3 | 14 | 11 | 15 |
| RS | Deltamethrin | 43 | 1 | 9 | 15 | 18 |
| RR | Untreated | 70 | 0 | 40 | 0 | 30 |
| RR | Permethrin | 53 | 0 | 41 | 0 | 12 |
| RR | Deltamethrin | 50 | 4 | 12 | 12 | 22 |

SS: homozygous susceptible genotype, RS: heterozygous genotype, RR: homozygous resistant genotype, KD: Knockdown.

### Impact of the *kdr* mutation on feeding success and biting behaviour

When female mosquitoes were pre-exposed to UTN (i.e., untreated netting), no difference in their feeding success was found among the three genotypes (OR_RS-SS_ = 1.47 [0.67, 3.22]; OR_RR-SS_ = 1.47 [0.68, 3.19]; OR_RR-RS_ = 1.00 [0.44, 2.25]; Figure 2A).

**Figure 2.**
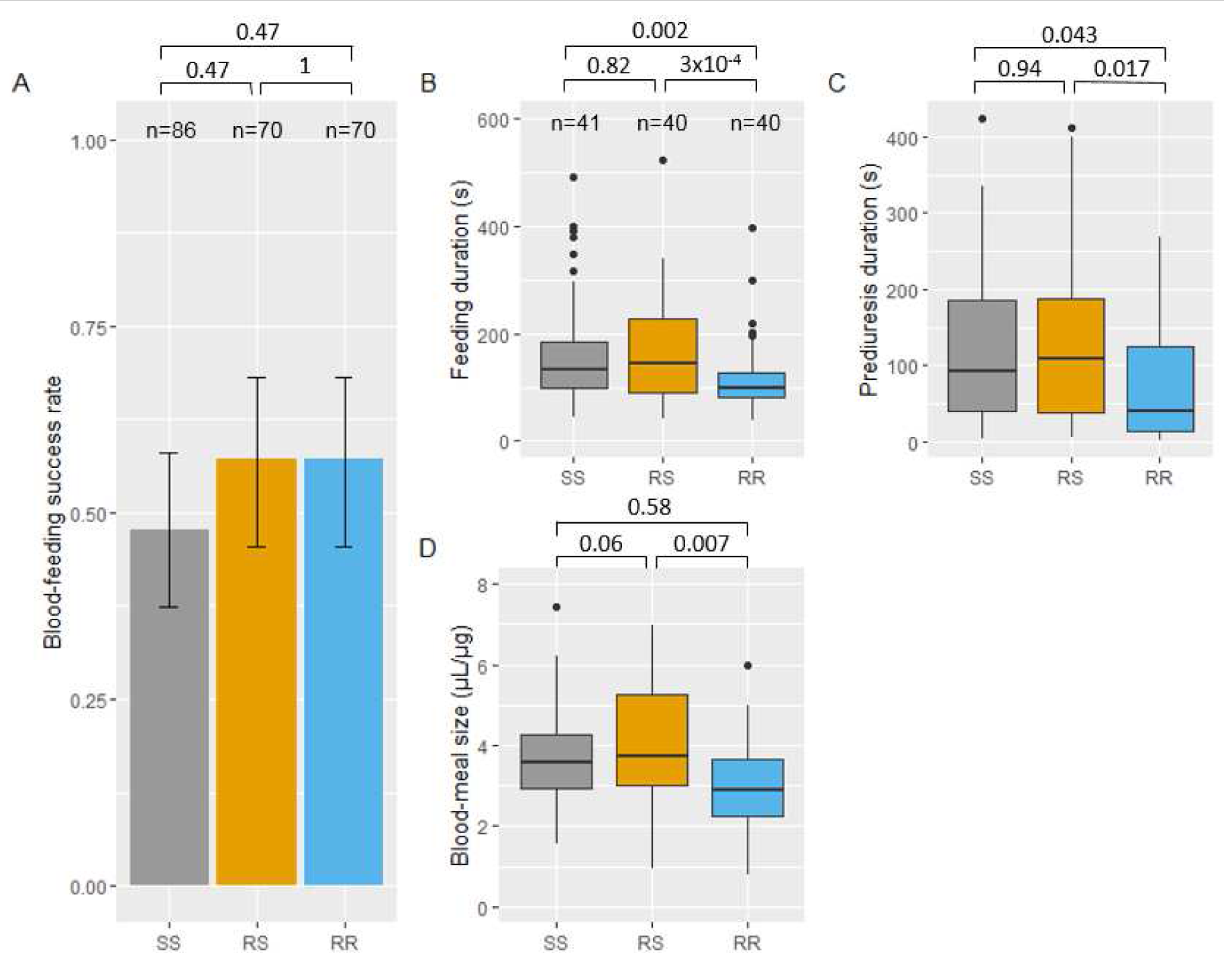
Feeding success and biting behavior in absence of insecticide of *Anopheles gambiae* females of each *kdr* genotype. Feeding success of each genotype and 95% binomial confidence intervals of the proportions (error bars) are shown in panel A (numbers n of mosquitoes exposed to the untreated net are indicated). Panels B, C and D show boxes-and-whiskers plots of feeding duration, prediuresis duration and blood-meal size, respectively (numbers n of blood-fed mosquitoes tested for behavioural parameters are indicated in panel B). Boxes indicate 1st-3rd quartile and median values. Whiskers indicate 1.5 inter-quartile range. P-values according to Tukey’s test after binomial mixed-effect model (panel A), mixed-effect cox proportional hazard model (panels B and C) and after linear mixed effect model (panel D) are indicated.

Analysis of the biting behaviour of successful mosquitoes in absence of insecticide pre-exposure showed that feeding and prediuresis durations were shorter in RR than in both SS and RS mosquitoes (feeding duration: HR_RR-SS_ = 2.15 [1.26, 3.69]; HR_RR-RS_ = 2.46 [1.43, 4.23]; Figure 2B; prediuresis duration: HR_RR-SS_ = 1.94 [1.02, 3.71]; HR_RR-RS_ = 2.11 [1.11, 4.01]; Figure 2C). The weighted blood meal volume of RR mosquitoes was lower than that of RS mosquitoes (MD_RS-RR_ = 0.99 [0.24, 1.74]; Figure 2D)). Number of probing events and probing duration were not significantly different among genotypes (SS, RS and RR) (Supplementary Tables 1 and 2).

### Impact of insecticide exposure on knockdown rates

Exposure to permethrin or deltamethrin induced 76 % [62.6, 85.7] and 41.3% [28.3, 55.7] KD rates in SS mosquitoes, respectively. It induced 32.6% [20.5, 47.5] and 37.2% [24.4, 52.1] KD rates in RS mosquitoes, respectively (Table 1). Among RR mosquitoes, permethrin exposure did not induce any KD effect, whereas deltamethrin exposure led to 30.2% [20.8, 45.8] of KD mosquitoes (Table 1). The *kdr* genotype was highly correlated with KD rates of mosquitoes exposed to permethrin (OR_RR-RS_ = 0.019 [6.1×10-4, 0.597], OR_RR-SS_ = 0.003 [9.7×10-5, 0.095], OR_RS-SS_ = 0.160 [5.41×10-2, 0.471]) but not for mosquitoes exposed to deltamethrin (OR_RR-RS_ = 0.80 [0.29, 2.22], OR_RR-SS_ = 0.67 [0.25, 1.83], OR_RS-SS_ = 0.85 [0.31, 2.34]).

### Impact of insecticide exposure on feeding success

When compared to the UTN condition, exposure to permethrin reduced significantly the feeding success of SS mosquitoes (OR_perm-UTN_ = 0.28 [0.10, 0.73]; Figure 3A), but not that of RS (OR_perm-UTN_ = 0.49 [0.19, 1.26]; Figure 3B). Feeding success of RR mosquitoes was higher (marginally not significant) (OR_perm-UTN_ = 2.58 [0.97, 6.84]; Figure 3B) when exposed to permethrin. Exposure to deltamethrin reduced significantly the feeding success of RS and RR mosquitoes (OR_delta-UTN_ = 0.23 [0.08, 0.64] and OR_delta-UTN_ = 0.35 [0.14 – 0.88]; Figure 3B and 3C), but not of SS mosquitoes (OR_delta-UTN_ = 0.58 [0.24, 1.43]; Figure 3A). When excluding KD mosquitoes from this analysis, all trends were kept (supplementary Figure 1) except that feeding success of SS mosquitoes was no longer reduced by permethrin exposure when compared to UTN (OR_perm-UTN_ = 0.78 [0.18, 3.40]; supplementary Figure 1A). Indeed among SS mosquitoes, non-KD had higher feeding success than those recorded as KD (OR_nonKD-KD_ = 4.71 [1.06, 20.9]; supplementary Table 3). The same was true for SS mosquitoes exposed to deltamethrin (OR_nonKD-KD_ = 22.5 [2.59, 195]; supplementary Table 3).

**Figure 3.**
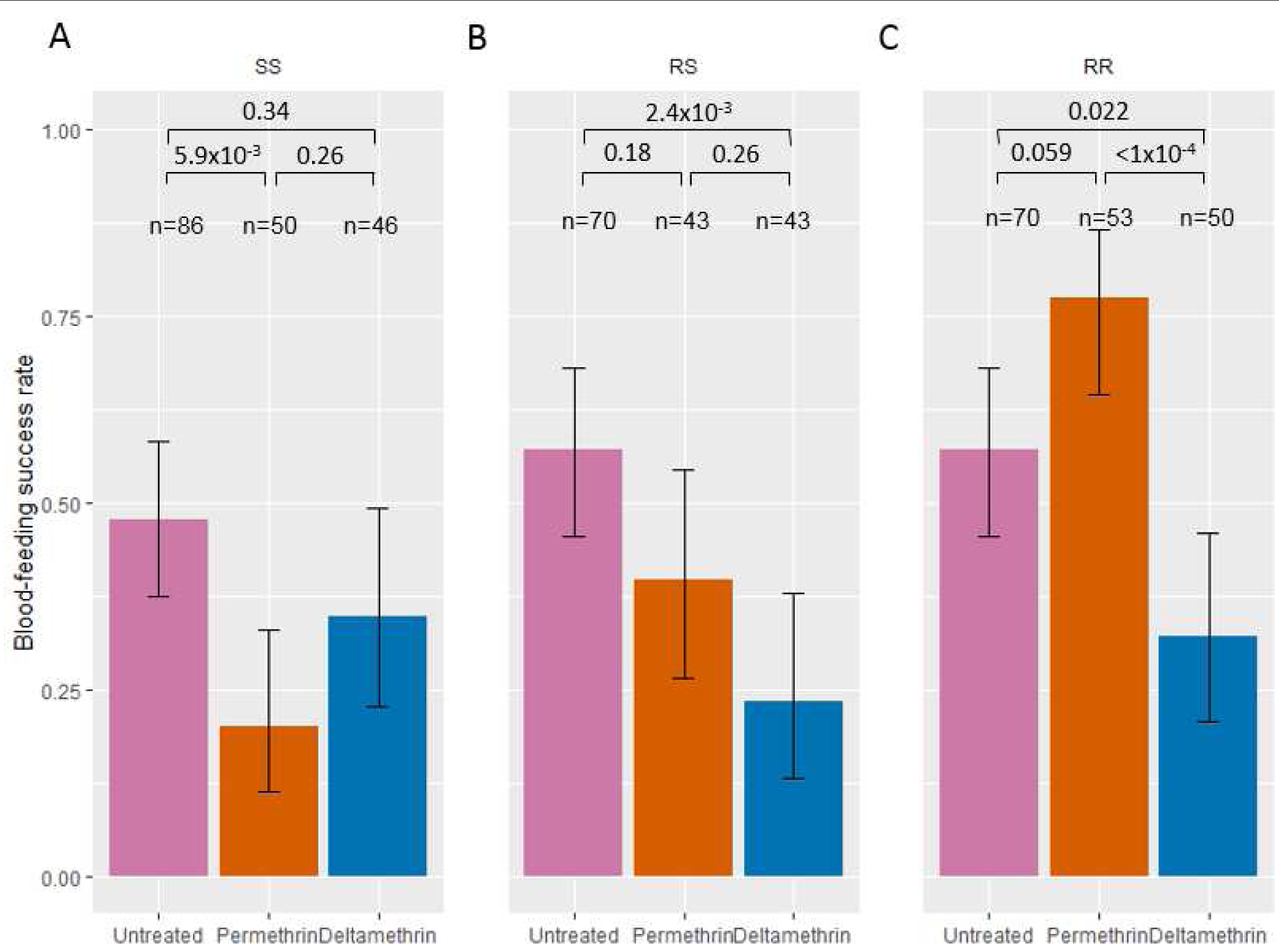
Feeding success after exposure to insecticides of *Anopheles gambiae* females of each *kdr* genotype. Feeding success of SS, RS, and RR (panels A, B and C, respectively) genotypes when exposed to untreated, permethrin-treated (Olyset) and deltamethrin-treated (PermaNet) nettings are shown with 95% binomial confidence intervals of the proportions (error bars). Numbers n of mosquitoes exposed to each treatment and for each genotype are indicated. P-values according to Tukey’s test after binomial mixed-effect model is indicated.

When comparing the feeding success among genotypes after insecticide exposure, the feeding rate of RR mosquitoes was higher than that of SS and RS mosquitoes after permethrin exposure (OR_RR-SS_ = 13.82 [4.35, 43.92]; OR_RR-RS_ = 5.29 [1.76, 15.86], Supplementary Figure 1B), whereas the feeding success of RS mosquitoes was although higher, not significantly different than that of SS mosquitoes (OR_RS-SS_ = 2.61 [0.85, 8.02], Supplementary Figure 2B). In contrast, exposure to deltamethrin did not induce any difference in the feeding success of the three genotypes (OR_RR-SS_ = 0.891 [0.31, 2.53]; OR_RR-RS_ = 1.55 [0.51, 4.76]; OR_RS-SS_ = 0.57 [0.18, 1.80], Supplementary Figure 2C). We observed the same trends when excluding KD mosquitoes from the analysis (supplementary Figure 3).

### Impact of insecticide exposure on biting behaviour

After exposure to deltamethrin, feeding duration, prediuresis duration and weighted blood meal size were significantly reduced in SS mosquitoes compared to the UTN condition (feeding duration: HR_delta-UTN_ = 4.38 [2.09, 9.15]; prediuresis: HR_delta-UTN_ = 5.31 [2.11, 13.36]; blood meal size; MD_delta-UTN_ = −1.02 [−1.82, −0.23]; Figure 4A, 4B and 4C, respectively). A similar trend was observed after exposure to permethrin for feeding duration (HR_perm-UTN_ = 2.87 [1.21, 6.81]; Figure 4A) and weighted blood meal volume (MD_perm-UTN_ = −0.97 [−1.85, −0.08]; Figure 4C) but not for prediuresis duration (HR_perm-UTN_ = 2.27 [0.92, 5.63]; Figure 4B). For this latter parameter, the non-significance was probably due to a lack of power.

**Figure 4.**
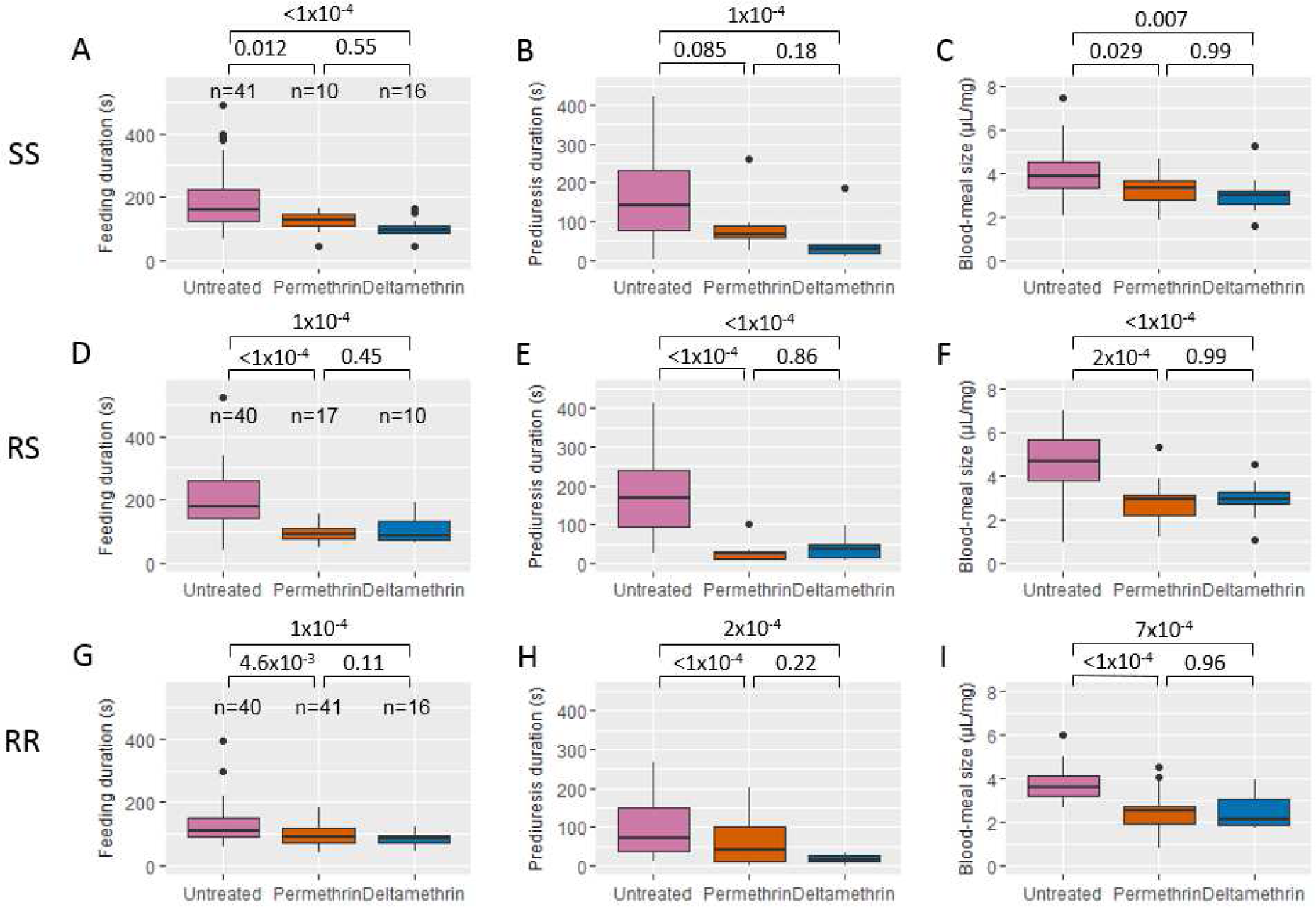
Biting behavior after exposure to insecticides of *Anopheles gambiae* females of each *kdr* genotype. Boxes-and-whiskers plots of feeding duration (panels A, D and M), prediuresis duration (panels B, E and H) and blood-meal size (panels C, F and I) are shown for each genotypes SS (panels A, B and C), RS (panels, D, E and F) and RR (panels M, H and I). Boxes indicate 1st-3rd quartile and median values. Whiskers indicate 1.5 inter-quartile range. Numbers n of blood-fed mosquitoes for each genotypes and tested for behavioural parameters are indicated in panels A, D and M. P-values according to Tukey’s test after mixed-effect cox proportional hazard model (panels A, B, D, E, M and H) and after linear mixed effect model (panels D, F and I) are indicated.

Compared to UTN, permethrin and deltamethrin reduced the feeding and prediuresis durations as well as blood meal size of both RS (feeding duration: HR perm-UTN = 7.42 [3.55, 15.53], HR_delta-UTN_ = 4.57 [1.94, 10.76], Figure 4D; prediuresis duration: HR_perm-UTN_ = 9.44 [3.36, 26.46] and HR_delta-UTN_ = 7.22 [2.70, 19.29], Figure 4E, blood-meal size: MD_perm-UTN_ = −1.97 [−3.04, −0.90] and MD_delta-UTN_ = −2.01 [−2.97, −1.05], Figure 4F) and RR genotypes (feeding duration: HR_perm-UTN_ = 2.06 [1.20, 3.51] and HR_delta-UTN_ = 3.78 [1.83, 7.82], Figure 4M; prediuresis duration: HR_perm-UTN_ = 1.71 [0.80, 3.64] and HR_delta-UTN_ = 9.31 [3.57, 24.28], Figure 4H; blood-meal size: MD_perm-UTN_ = −1.52 [−2.23, −0.81] and MD_delta-UTN_ = −1.41 [−2.28, −0.54], Figure 4I). Number of probing events and probing duration were not significantly different among treatments (UTN and ITNs) whatever the genotype (Supplementary Tables 4 and 5).

When comparing the biting behavior among genotypes after insecticide exposure, we found that prediuresis duration of RS mosquitoes was shorter than that of SS mosquitoes after permethrin exposure (HR_RS-_ SS = 3.82 [1.15, 12.7], supplementary Table 8). Moreover, prediuresis duration of RR mosquitoes was shorter than that of SS after deltamethrin exposure (HR_RR-SS_ = 3.41 [1.13, 10.29], supplementary Table 8). For all other parameters, we were not able to evidence any differences among genotypes (Supplementary Tables 1, 2, 7 and 8).

In absence of exposure to insecticide (UTN), the blood meal size and the prediuresis duration were positively correlated with the feeding duration for all genotypes (supplementary Tables 6 and 7). With the exception of blood meal size of RS exposed to permethrin, these relationships were not observed when mosquitoes were exposed to insecticides (supplementary Tables 9 and 10), suggesting a perturbation of processes underlying these correlations.

Excluding 14 KD mosquitoes from the analyses of the biting behaviour parameters do not significantly change the results (supplementary Tables 11 to 22).

## Discussion

To investigate the influence of ITN exposure on the biting behaviour of *An. mambiae* mosquitoes, we used mosquitoes that share the same genetic background, but for the *kdr* allele locus, and then exposed them to ITN prior to blood feeding.

The blood-feeding success did not differ between the three genotypes in the absence of insecticide exposure. Therefore, the *kdr* mutation was not associated with significant change of blood meal success rate. However, feeding duration and blood meal size were different between genotypes. RR mosquitoes spent less time taking their blood meal than RS and SS mosquitoes. This might confer an advantage as fast feeding reduces the risk to be killed because of the host defensive behaviour (48). On the other hand, RS mosquitoes took higher blood volumes than RR females. This could improve the completion of oogenesis in RS mosquitoes (49) and increase their fecundity compared to RR (50). However, large blood meals reduce the flying ability, escape speed and agility required to avoid predators (48,51,52). These different trades-offs between behavioural traits that might enhance fecundity or survival in the three genotypes are of great interest and deserve further investigations in relation with the ecological and vector control environment. Such trades-offs possibly affect mosquito fitness and may therefore drive not only the evolution of insecticide resistance in mosquitoes but also parasite transmission. For example, a decrease in blood meal duration and size might increase the frequency of multiple feedings and consequently the risk of *Plasmodium* transmission (53). Similarly, a bigger blood meal size might increase the probability of mosquito infection by gametocytes (54).

Exposure to permethrin and deltamethrin induced opposite outcomes in term of blood feeding success (increase and decrease, respectively) in RR mosquitoes. This opposite effect on the feeding success rate of RR females might be linked to the different chemical properties of permethrin and deltamethrin that induce two types of bursting activity of sodium channels (55,56). Type II pyrethroids, such as deltamethrin, further delay the inactivation of the voltage-gated sodium channel and in a less reversible way than type I pyrethroids, such as permethrin (57). The lower PYR susceptibility of homozygous resistant mosquitoes could lead to their over-stimulation compared to susceptible and heterozygous mosquitoes that are more affected by the toxic effect of such insecticides (58).

In contrast, among females that have been successful in taking a blood meal, the behavioural sequence was altered in the same way by both insecticides. They both induced a decrease of feeding duration and prediuresis duration. This is in agreement with the results of Hauser *et al*. (50) showing that mosquitoes biting trough permethrin+PBO nets (Olyset Plus) had less feeding successes, shorter feeding duration and lower blood meal sizes compared to those biting trough untreated nets. As discussed above, short feeding durations and small blood-meal lead us to expect that exposure to insecticide may (i) reduce the risk of the vector to be killed due to the defensive behaviour of the host or due to predators, (ii) reduce the possible number of parasites ingested during one feeding attempt that may be compensated by (iii) the increase of multiple feeding (and therefore the higher risk of human-to-mosquito and mosquito-to-human parasite transmission).

Whatever the genotype, blood meal size and prediuresis duration were correlated to feeding duration in absence of exposure to insecticides. This relationship was, in most cases, no longer observed when mosquitoes were exposed to insecticides indicating that sub-lethal contact with insecticides disrupt physiological processes involved in blood meal intake. Such indirect evidence highlight the need to further investigate the consequences of sub-lethal contact with insecticides.

Prediuresis duration was substantially reduced in RS and RR females after exposure to ITNs. Prediuresis is an intestinal mechanism that plays a crucial role in protein concentration during feeding (59) and contributes to thermoregulation (60,61). This perturbation of the prediuresis phase by sub-lethal doses of PYR insecticides could lead to toxic accumulation of metabolic wastes and products of oxidative stress in the haemolymph that might affect the lifespan of mosquitoes. This results suggest that mosquitoes with the *kdr* mutation might be more susceptible to new chemicals that target the mosquito renal system and that are currently developed as an alternative to the currently used insecticides (62).

As expected, we found a strong relationship between *kdr* genotype and KD phenotype when mosquitoes were exposed to Permethrin. However with deltamethrin, we were not able to find such a relationship and mosquitoes carrying the *kdr* mutation experienced moderate levels of KD. Indeed, deltamethrin is expected to induce higher knock-down rates than permethrin against resistant populations of *An. mambiae* (63). This observation is also true when looking at mortality in experimental hut trial in areas with high frequencies of the *kdr* mutation in the vector population (64–66). This difference between permethrin and deltamethrin effect may be linked to the different chemical properties of permethrin and deltamethrin (type I and Type II pyrethroids) as describe above.

We found that KD reduced the feeding success of mosquitoes exposed to PYR insecticides, particularly in SS mosquitoes. However when analyzing feeding success and behavior of non-KD mosquitoes carrying the *kdr* mutation, we observed the same trends than we get when including KD mosquitoes. This indicates that observed differences in feeding success and behavior are therefore directly linked to the presence of the mutation and not only a consequence of the KD phenotype.

This work has some limitations. First, we were not able to randomize genotypes over time because of rearing constraints. Consequently all RS mosquitoes were tested during the last month of experiments. This did not induce any effect when analyzing the treatment effect relative to the genotype but might have introduced a bias while comparing RS with RR and/or SS mosquitoes (as experimental period is possibly a confounding factor for RS genotype). The second limitation relies on body size measurement. Indeed, we chose to use an easy method to get a proxy of mosquito size. We randomly selected five anopheles females from each rearing cages used during the experiment, weighed them together and used the mean weight to adjust blood-meal size. An individual wing length measurement would allow to avoid any bias in developmental variability within each rearing cage.

To conclude, our study demonstrates a complex interaction between insecticide exposure and the *kdr* mutation on the biting behaviour of mosquitoes. The behavioural modulation induced by PYR-treated nets also raises concerns about the consequences of the *kdr* resistance-insecticide interaction. In previous studies, we evidenced that RR mosquitoes prefer a host protected by a permethrin-treated net rather than an untreated net (67) and that heterozygotes RS mosquitoes have a remarkable ability to find a hole into a bet net (25). Herein, we have completed the sequence by showing that *kdr* homozygous resistant *An. mambiae* displayed enhanced feeding success when exposed to permethrin ITN. However here, we exposed all our mosquitoes for a constant duration to the treated nets which does not reflect the variability that happens in natural conditions. Indeed, we can expect from the literature (22,23,25) that both the genotype for the *kdr* mutation and the type of pyrethroids on LLINs may affect the contact duration. It would be of great interest to decipher with the relationship between the time of contact with the insecticide and the feeding sequence.

Insecticide resistance genes in malaria vectors could modify vector competence and the dynamics of infection by *P. falciparum*. For instance, recent studies have shown that parasite infection increases insecticide susceptibility in mosquitoes carrying the *kdr* mutation (68) and that insecticide exposure reduces parasite development in resistant mosquitoes (69). In addition, malaria parasites have been shown to modify the feeding behaviour of their mosquito vectors in ways that favour their transmission (70–72), but the role of insecticide resistance that could modulate this phenomenon has not been investigated yet (73). It is therefore urgent to decipher the links between insecticide exposure, resistance mechanisms and infection by *P. falciparum* on the host-seeking and biting behaviour of malaria vectors to better understand malaria transmission in areas where insecticidal tools for malaria prevention are implemented. All these interactions should then be used as variables to include host-seeking behavioural modulation by *kdr* resistance in models of resistance evolution and *P. falciparum* transmission to better understand and/or predict the efficacy of vector control strategies (74).

## Data accessibility

Data are available online: doi:10.5281/zenodo.2616599

## Supplementary material

Script and codes are available online: doi:10.5281/zenodo.2616599

Supplementary Tables and Figures are available online: doi:10.5281/zenodo.3901471

## Acknowledgements

MD was supported by IRD’s PhD student fellowship. The authors would like to acknowledge the technical/research platform dedicated on vectors at the IRD centre and all members of the platform for providing mosquitoes and technical support. This platform is a member of the Vectopole Sud network and of the LabEX CEMEB (Centre Méditerranéen de l’Environnement et de la Biodiversité) in Montpellier. We thank Dr Anna Cohuet and Dr Thierry Lefèvre for their valuable comments on the manuscript. We also thank Dr Etienne Bilgo, Dr Niels Verhulst, Dr Thomas Muillemaud and one anonymous reviewer for their relevant comments which helped improve this manuscript. Version 5 of this preprint has been peer-reviewed and recommended by Peer Community In Zool (https://doi.org/10.24072/pci.zool.100003).

## Conflict of interest disclosure

The authors of this preprint declare that they have no financial conflict of interest with the content of this article. NM and CP are recommenders for PCI Zool.

## Notes

http://doi.org/10.5281/zenodo.3629451

http://doi.org/doi:10.5281/zenodo.3901471

### Competing Interest Statement

The authors have declared no competing interest.

### Summary of Updates

Version 5 of this preprint has been peer-reviewed and recommended by Peer Community In Zool (https://doi.org/10.24072/pci.zool.100003). The name of one reviewer has been added.

