## Supplementary Figures and Tables for "Sub-lethal insecticide exposure affects host biting efficiency of *Kdr*-resistant *Anopheles gambiae*"

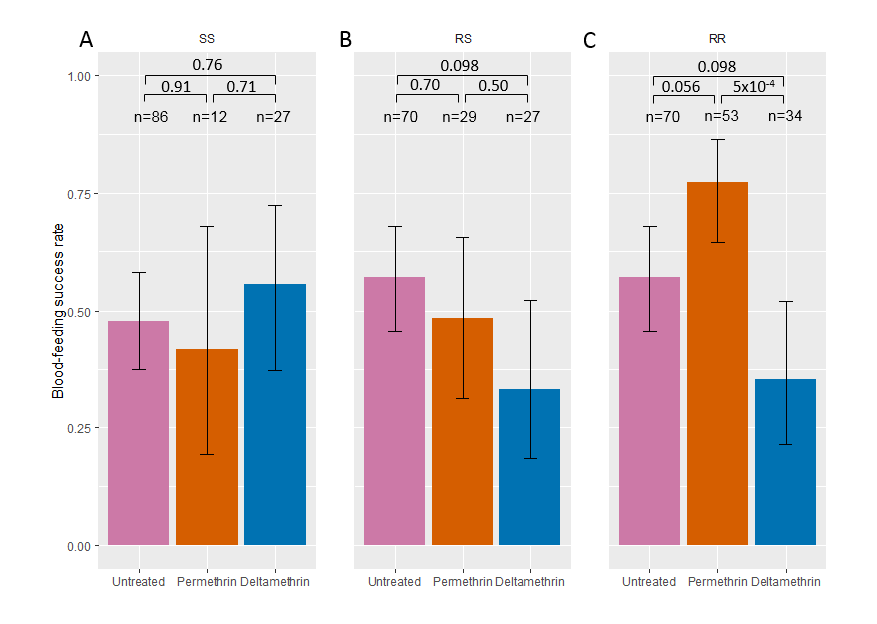

**Supplementary Figure 1. Feeding success after exposure to insecticides of non-KD *Anopheles gambiae* females of each *kdr* genotype.** Feeding success of non-KD SS, RS, and RR (panels A, B and C, respectively) genotypes when exposed to untreated, permethrin-treated (Olyset) and deltamethrin-treated (PermaNet) nettings are shown with 95% binomial confidence intervals of the proportions (error bars). Numbers n of mosquitoes exposed to each treatment and for each genotype are indicated. P-values according to Tukey’s test after binomial mixed-effect model is indicated.

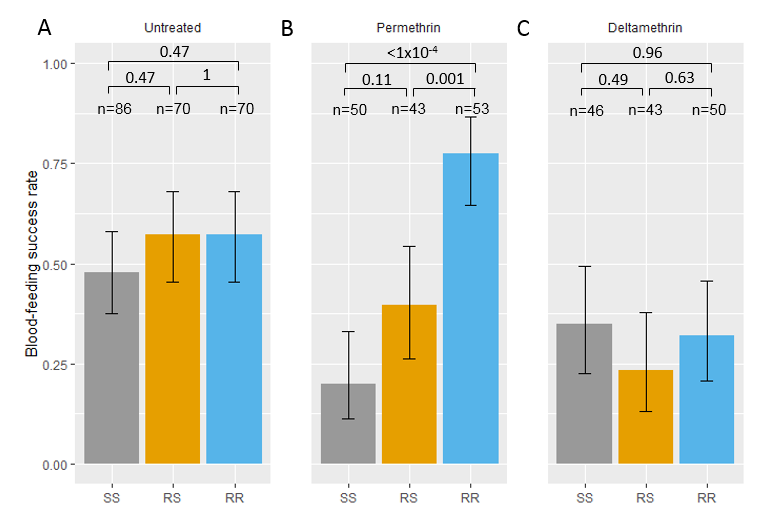

Supplementary Figure 2. Among *kdr* genotypes comparison of feeding success of *Anopheles gambiae* females after exposure to insecticides. Feeding success of each genotype and 95% binomial confidence intervals of the proportions (error bars) are shown after exposure to untreated net (Panel A, same as Figure 2A), to permethrin treated net (Olyset; panel B) and to deltamethrin treated net (Permanet; panel C). Numbers n of mosquitoes exposed to each treatment and for each genotype are indicated. Significance according to Tukey’s test after binomial mixed-effect model is indicated.

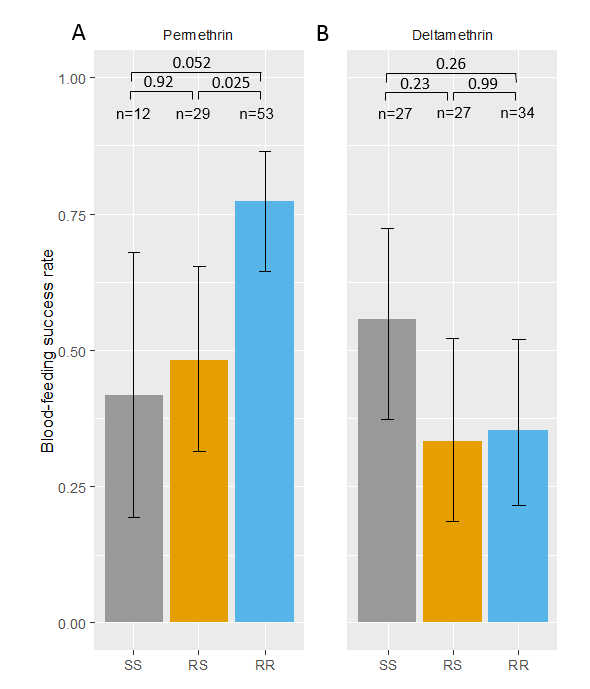

Supplementary Figure 3. Among kdr genotypes comparison of feeding success of non-KD Anopheles gambiae females after exposure to insecticides. Feeding success of non-KD mosquitoes of each genotype and 95% binomial confidence intervals of the proportions (error bars) are shown after exposure to permethrin treated net (Olyset; panel A) and to deltamethrin treated net (Permanet; panel B). Numbers n of mosquitoes exposed to each treatment and for each genotype are indicated. Significance according to Tukey’s test after binomial mixed-effect model is indicated.

Supplementary Table 1: Among genotype comparison of the number of probing events of Anopheles mosquitoes

| Treatment | contrast | Rate-Ratio (95% CI) | p-value |
| --- | --- | --- | --- |
| Permanet | RR / RS | 0.2065269 [0.01982615, 2.151369] | 0.25282895 |
|  | RR / SS | 4.3650588 [0.20049661, 95.032722] | 0.49737134 |
|  | RS / SS | 21.1355422 [0.93350901, 478.529016] | 0.05681498 |
| Olyset | RR / RS | 0.1996279 [0.02935438, 1.357593] | 0.11870444 |
|  | RR / SS | 0.3023033 [0.03268575, 2.795936] | 0.41419506 |
|  | RS / SS | 1.5143334 [0.147877, 15.507522] | 0.90700469 |
| Untreated Net | RR / RS | 1.4266078 [0.3367924, 6.042921] | 0.83053756 |
|  | RR / SS | 0.7093201 [0.19364229, 2.59827] | 0.80697994 |
|  | RS / SS | 0.4972075 [0.11593091, 2.132436] | 0.49514241 |

Supplementary Table 2: Among genotype comparison of the hazard to complete probing of Anopheles mosquitoes

| Treatment | contrast | Hazard-Ratio (95% CI) | p-value |
| --- | --- | --- | --- |
| Permanet | RR / RS | 1.5899262 [0.6135174, 4.120283] | 0.4886324 |
|  | RR / SS | 0.9727067 [0.423672, 2.233233] | 0.9966484 |
|  | RS / SS | 0.6117936 [0.2349056, 1.593369] | 0.4513116 |
| Olyset | RR / RS | 1.799743 [0.9046358, 3.580529] | 0.1116432 |
|  | RR / SS | 0.939998 [0.4107093, 2.151391] | 0.98323 |
|  | RS / SS | 0.5222957 [0.20268, 1.345929] | 0.2421304 |
| Untreated Net | RR / RS | 1.0652724 [0.6267428, 1.810639] | 0.9578886 |
|  | RR / SS | 0.9760263 [0.5737765, 1.660276] | 0.9937017 |
|  | RS / SS | 0.9162223 [0.5396132, 1.555676] | 0.9206321 |

Supplementary Table 3: Comparison of feeding success between non-KD and KD mosquitoes

| Treatment | genotype | Odds-Ratio [95% CI] | p-value |
| --- | --- | --- | --- |
| Permethrin | RR | 1.6363496 [0.4291883, 6.238847] | 0.46942047 |
|  | RS | 7.5003577 [0.8424351, 66.777095] | 0.07072763 |
|  | SS | 22.499333 [2.590607, 195.405971] | 0.0049162 |
| Deltamethrin | RR | 0.7482277 [2.220446E-16, Inf] | 0.99999982 |
|  | RS | 3.4221854 [0.7819664, 14.976799] | 0.10200971 |
|  | SS | 4.7143423 [1.062193, 20.923708] | 0.04147763 |

Supplementary Table 4: Among treatment comparison of the number of probing events for each genotype

| genotype | contrast | Rate-Ratio (95% CI) | p-value |
| --- | --- | --- | --- |
| RR | Deltamethrin / Permethrin | 2.06917474 [0.260680975, 16.424229] | 0.6858793 |
|  | Deltamethrin / Untreated | 0.61063592 [0.093900111, 3.970988] | 0.808396 |
|  | Permethrin / Untreated | 0.29511085 [0.060249141, 1.445505] | 0.1678518 |
| RS | Deltamethrin / Permethrin | 2.00005422 [0.221198567, 18.08428] | 0.7382395 |
|  | Deltamethrin / Untreated | 4.21803553 [0.552322234, 32.21276] | 0.2187943 |
|  | Permethrin / Untreated | 2.10896059 [0.32616651, 13.636332] | 0.6134566 |
| SS | Deltamethrin / Permethrin | 0.14330122 [0.006023182, 3.409367] | 0.318958 |
|  | Deltamethrin / Untreated | 0.09922806 [0.006276947, 1.56863] | 0.1208565 |
|  | Permethrin / Untreated | 0.69244396 [0.092505398, 5.18325] | 0.9027997 |

Supplementary Table 5: Among treatment comparison of the hazard to complete probing for each genotype

| genotype | contrast | Hazard-Ratio (95% CI) | p-value |
| --- | --- | --- | --- |
| RR | Deltamethrin / Permethrin | 0.969668 [0.4851137, 1.938218] | 0.994028 |
|  | Deltamethrin / Untreated | 1.0944956 [0.5441664, 2.201386] | 0.9507035 |
|  | Permethrin / Untreated | 1.1287323 [0.6660412, 1.912849] | 0.8525701 |
| RS | Deltamethrin / Permethrin | 1.0976316 [0.4294804, 2.805239] | 0.9705967 |
|  | Deltamethrin / Untreated | 0.7333271 [0.3189066, 1.686289] | 0.657375 |
|  | Permethrin / Untreated | 0.6680994 [0.3370059, 1.324478] | 0.350725 |
| SS | Deltamethrin / Permethrin | 0.9370615 [0.3638974, 2.413] | 0.9857986 |
|  | Deltamethrin / Untreated | 1.0982308 [0.5479872, 2.200984] | 0.9464827 |
|  | Permethrin / Untreated | 1.1719944 [0.5103472, 2.691444] | 0.8955522 |

Supplementary Table 6: Among treatments comparison of weighted volume of blood meal

| treatment | contrast | Difference [95% CI] | p-value |
| --- | --- | --- | --- |
| Permanet | RR / RS | -0.3889038 [-1.51541812, 0.7376106] | 0.6894645 |
|  | RR / SS | -0.6642439 [-1.6975565, 0.3690687] | 0.28036573 |
|  | RS / SS | -0.2753401 [-1.38489115, 0.8342109] | 0.82553138 |
| Olyset | RR / RS | -0.5383489 [-1.59096245, 0.5142646] | 0.43324541 |
|  | RR / SS | -0.8318678 [-1.79150699, 0.1277714] | 0.10355568 |
|  | RS / SS | -0.2935189 [-1.49418704, 0.9071493] | 0.82894899 |
| Untreated Net | RR / RS | -0.9912139 [-1.744629, -0.2377987] | 0.00715823 |
|  | RR / SS | -0.2814588 [-0.95281868, 0.389901] | 0.57635249 |
|  | RS / SS | 0.709755 [-0.02932594, 1.448836] | 0.06220632 |

Supplementary Table 7: Among genotype comparison of the hazard to complete feeding of Anopheles mosquitoes

| treatment | contrast | Hazard ratio [95% CI] | p-value |
| --- | --- | --- | --- |
| Permanet | RR / RS | 2.0351811 [0.7774306, 5.327758] | 0.19376895 |
|  | RR / SS | 1.8595024 [0.8048335, 4.296229] | 0.19177081 |
|  | RS / SS | 0.9136791 [0.3532432, 2.363271] | 0.97304429 |
| Olyset | RR / RS | 0.6810666 [0.3437089, 1.349548] | 0.38600539 |
|  | RR / SS | 1.5405116 [0.6718357, 3.532375] | 0.44105525 |
|  | RS / SS | 2.2619102 [0.8805691, 5.810149] | 0.10558688 |
| Untreated Net | RR / RS | 2.4606925 [1.4299026, 4.234559] | 0.00029811 |
|  | RR / SS | 2.1527285 [1.2571208, 3.686392] | 0.00240853 |
|  | RS / SS | 0.8748466 [0.5137238, 1.489821] | 0.8262213 |

Supplementary Table 8: Among genotype comparison of the hazard to complete prediuresis of Anopheles mosquitoes

| treatment | contrast | Hazard ratio [95% CI] | p-value |
| --- | --- | --- | --- |
| Permanet | RR / RS | 2.7197592 [0.8761514, 8.442708] | 0.09608853 |
|  | RR / SS | 3.4080335 [1.1292817, 10.285026] | 0.02514575 |
|  | RS / SS | 1.2530644 [0.396447, 3.960606] | 0.89018754 |
| Olyset | RR / RS | 0.382092 [0.1295448, 1.126979] | 0.09299368 |
|  | RR / SS | 1.4608704 [0.5456537, 3.911166] | 0.63900287 |
|  | RS / SS | 3.8233472 [1.1514099, 12.695725] | 0.02395076 |
| Control | RR / RS | 2.1089395 [1.1092902, 4.009434] | 0.01782237 |
|  | RR / SS | 1.9427912 [1.0175798, 3.70923] | 0.04252618 |
|  | RS / SS | 0.9212171 [0.5104568, 1.662513] | 0.94318562 |

Supplementary Table 9: Marginal slopes of the linear trend between feeding duration and blood-meal size.

| genotype | treatment | Marginal slope | |
| --- | --- | --- | --- |
|  |  | Value | 95% confidence interval |
| SS | Untreated | 0.003007 | [0.000442, 0.00557] |
| RS | Untreated | 0.003964 | [0.00119, 0.00674] |
| RR | Untreated | 0.005862 | [0.00167, 0.01005] |
| SS | Permethrin | 0.009937 | [-0.008062, 0.02794] |
| RS | Permethrin | 0.023022 | [0.00776, 0.03828] |
| RR | Permethrin | 0.004893 | [-0.002253, 0.01204] |
| SS | Deltamethrin | 0.000355 | [-0.012805, 0.01352] |
| RS | Deltamethrin | 0.005265 | [-0.006496, 0.01703] |
| RR | Deltamethrin | 0.01763 | [-0.006509, 0.04177] |

Supplementary Table 10: Marginal slope of the linear trend between feeding duration and the hazard of completing pre-diuresis.

| genotype | treatment | Marginal slope | |
| --- | --- | --- | --- |
|  |  | Value | 95% confidence interval |
| SS | Untreated | -0.00629 | [-0.00904, -0.00354] |
| RS | Untreated | -0.00302 | [-0.00496, -0.00107] |
| RR | Untreated | -0.00616 | [-0.01095, -0.00137] |
| SS | Permethrin | -0.01087 | [-0.02967, 0.00793] |
| RS | Permethrin | 0.75192 | [-0.3316, 1.83544] |
| RR | Permethrin | -0.00836 | [-0.02272, 0.00599] |
| SS | Deltamethrin | 0.31275 | [-0.02684, 0.65235] |
| RS | Deltamethrin | -0.02304 | [-0.04829, 0.0022] |
| RR | Deltamethrin | 0.82761 | [-0.58179, 2.23701] |

Supplementary Table 11: Among treatments comparison of feeding success for non-KD mosquitoes

| genotype | contrast | Odds-Ratio [95% CI] | p-value |
| --- | --- | --- | --- |
| RR | Deltamethrin / Permethrin | 0.16 [0.0508, 0.501] | 0.0005 |
|  | Deltamethrin / Untreated | 0.409 [0.1479, 1.132] | 0.0982 |
|  | Permethrin / Untreated | 2.562 [0.9825, 6.683] | 0.0557 |
| RS | Deltamethrin / Permethrin | 0.536 [0.1462, 1.963] | 0.4958 |
|  | Deltamethrin / Untreated | 0.375 [0.1229, 1.145] | 0.0979 |
|  | Permethrin / Untreated | 0.7 [0.2468, 1.986] | 0.7003 |
| SS | Deltamethrin / Permethrin | 1.75 [0.3356, 9.126] | 0.705 |
|  | Deltamethrin / Untreated | 1.372 [0.4834, 3.894] | 0.7558 |
|  | Permethrin / Untreated | 0.784 [0.1806, 3.403] | 0.9196 |

Supplementary Table 12: Among genotypes comparison of feeding success for non-KD mosquitoes

| Treatment | contrast | Odds-Ratio [95% CI] | p-value |
| --- | --- | --- | --- |
| Deltamethrin | RR / RS | 1.091 [0.304, 3.92] | 0.986 |
|  | RR / SS | 0.436 [0.126, 1.51] | 0.2596 |
|  | RS / SS | 0.4 [0.106, 1.5] | 0.2349 |
| Permethrin | RR / RS | 3.661 [1.14, 11.75] | 0.0249 |
|  | RR / SS | 4.783 [0.986, 23.2] | 0.0527 |
|  | RS / SS | 1.307 [0.256, 6.68] | 0.9213 |
| Untreated Net | RR / RS | 1 [0.448, 2.23] | 1 |
|  | RR / SS | 1.463 [0.683, 3.14] | 0.4686 |
|  | RS / SS | 1.463 [0.683, 3.14] | 0.4686 |

Supplementary Table 13: Among treatments comparison of number of probing events for non-KD mosquitoes

| genotype | contrast | Rate-Ratio [95% CI] | p-value |
| --- | --- | --- | --- |
| RR | Deltamethrin / Permethrin | 2.855 [0.33206, 24.54] | 0.4839 |
|  | Deltamethrin / Untreated | 0.84 [0.11796, 5.98] | 0.976 |
|  | Permethrin / Untreated | 0.294 [0.06034, 1.43] | 0.1645 |
| RS | Deltamethrin / Permethrin | 1.837 [0.18326, 18.41] | 0.8078 |
|  | Deltamethrin / Untreated | 4.804 [0.58837, 39.22] | 0.1842 |
|  | Permethrin / Untreated | 2.616 [0.38017, 18] | 0.4683 |
| SS | Deltamethrin / Permethrin | 0.32 [0.0061, 16.82] | 0.7764 |
|  | Deltamethrin / Untreated | 0.107 [0.00673, 1.69] | 0.1379 |
|  | Permethrin / Untreated | 0.333 [0.01489, 7.44] | 0.6812 |

Supplementary Table 14: Among genotypes comparison of number of probing events for non-KD mosquitoes

| treatment | contrast | Rate-Ratio [95% CI] | p-value |
| --- | --- | --- | --- |
| Permanet | RR / RS | 0.2478951 [0.02102436, 2.922894] | 0.37768122 |
|  | RR / SS | 5.6517541 [0.24348914, 131.185827] | 0.39664753 |
|  | RS / SS | 22.7989767 [0.95816029, 542.491005] | 0.05407603 |
| Olyset | RR / RS | 0.1594924 [0.02214653, 1.148615] | 0.07438331 |
|  | RR / SS | 0.6343896 [0.02444015, 16.466758] | 0.94177923 |
|  | RS / SS | 3.9775523 [0.13729325, 115.234523] | 0.59773377 |
| Untreated Net | RR / RS | 1.4180706 [0.33722814, 5.963097] | 0.8340559 |
|  | RR / SS | 0.7178176 [0.19751305, 2.60875] | 0.81669609 |
|  | RS / SS | 0.5061932 [0.1203679, 2.128736] | 0.50343206 |

Supplementary Table 15: Among treatments comparison of probing duration for non-KD mosquitoes

| genotype | contrast | Hazard-Ratio [95% CI] | p-value |
| --- | --- | --- | --- |
| RR | Deltamethrin / Permethrin | 0.808 [0.373, 1.75] | 0.793 |
|  | Deltamethrin / Untreated | 0.918 [0.422, 2] | 0.964 |
|  | Permethrin / Untreated | 1.137 [0.67, 1.93] | 0.8369 |
| RS | Deltamethrin / Permethrin | 1.168 [0.427, 3.19] | 0.9307 |
|  | Deltamethrin / Untreated | 0.659 [0.276, 1.57] | 0.498 |
|  | Permethrin / Untreated | 0.564 [0.27, 1.18] | 0.1612 |
| SS | Deltamethrin / Permethrin | 0.662 [0.197, 2.23] | 0.7052 |
|  | Deltamethrin / Untreated | 1.039 [0.51, 2.12] | 0.9911 |
|  | Permethrin / Untreated | 1.569 [0.514, 4.79] | 0.6112 |

Supplementary Table 16: Among genotype comparison of probing duration for non-KD mosquitoes

| treatment | contrast | Hazard-Ratio [95% CI] | p-value |
| --- | --- | --- | --- |
| Permanet | RR / RS | 1.4849422 [0.52457876, 4.203474] | 0.64628417 |
|  | RR / SS | 0.8601859 [0.34586729, 2.139317] | 0.92060356 |
|  | RS / SS | 0.5792723 [0.21268749, 1.577697] | 0.40811093 |
| Olyset | RR / RS | 2.1468983 [1.02502831, 4.496629] | 0.04087478 |
|  | RR / SS | 0.7054413 [0.23176682, 2.14719] | 0.74284818 |
|  | RS / SS | 0.3285863 [0.09540271, 1.131718] | 0.08795174 |
| Untreated Net | RR / RS | 1.0651534 [0.62613232, 1.812] | 0.95816734 |
|  | RR / SS | 0.9738627 [0.57206198, 1.657877] | 0.99252325 |
|  | RS / SS | 0.9142933 [0.53792485, 1.553995] | 0.91723635 |

Supplementary Table 17: Among treatments comparison of feeding duration for non-KD mosquitoes

| genotype | contrast | Hazard-Ratio [95% CI] | p-value |
| --- | --- | --- | --- |
| RR | Deltamethrin / Permethrin | 1.749 [0.8, 3.83] | 0.2151 |
|  | Deltamethrin / Untreated | 3.593 [1.61, 8.02] | 0.0005 |
|  | Permethrin / Untreated | 2.054 [1.202, 3.51] | 0.0047 |
| RS | Deltamethrin / Permethrin | 0.462 [0.166, 1.28] | 0.178 |
|  | Deltamethrin / Untreated | 4.189 [1.718, 10.21] | 0.0005 |
|  | Permethrin / Untreated | 9.072 [4.089, 20.13] | <.0001 |
| SS | Deltamethrin / Permethrin | 1.884 [0.56, 6.34] | 0.4392 |
|  | Deltamethrin / Untreated | 4.272 [2.013, 9.07] | <.0001 |
|  | Permethrin / Untreated | 2.267 [0.727, 7.07] | 0.21 |

Supplementary Table 18: Among genotypes comparison of feeding duration for non-KD mosquitoes

| treatment | contrast | Hazard-Ratio [95% CI] | p-value |
| --- | --- | --- | --- |
| Permanet | RR / RS | 2.1086406 [0.7374156, 6.02966] | 0.21907481 |
|  | RR / SS | 1.810375 [0.7239442, 4.527224] | 0.28254995 |
|  | RS / SS | 0.8585508 [0.3179152, 2.318572] | 0.9311368 |
| Olyset | RR / RS | 0.5566173 [0.2652585, 1.168003] | 0.15271709 |
|  | RR / SS | 1.9502887 [0.6407795, 5.935935] | 0.33747344 |
|  | RS / SS | 3.5038234 [1.0166501, 12.075717] | 0.04617139 |
| Untreated Net | RR / RS | 2.458579 [1.4284269, 4.231656] | 0.0003042 |
|  | RR / SS | 2.1527095 [1.256963, 3.68679] | 0.00241505 |
|  | RS / SS | 0.875591 [0.5141581, 1.491097] | 0.82822543 |

Supplementary Table 19: Among treatments comparison of feeding duration for non-KD mosquitoes

| genotype | contrast | Hazard-Ratio [95% CI] | p-value |
| --- | --- | --- | --- |
| RR | Deltamethrin / Permethrin | 6.02 [1.941, 18.65] | 0.0006 |
|  | Deltamethrin / Untreated | 10.38 [3.493, 30.86] | <.0001 |
|  | Permethrin / Untreated | 1.73 [0.81, 3.68] | 0.209 |
| RS | Deltamethrin / Permethrin | 1 [0.253, 3.96] | 1 |
|  | Deltamethrin / Untreated | 7.81 [2.776, 21.99] | <.0001 |
|  | Permethrin / Untreated | 7.81 [2.414, 25.25] | 0.0001 |
| SS | Deltamethrin / Permethrin | 1.83 [0.471, 7.13] | 0.5478 |
|  | Deltamethrin / Untreated | 5.19 [1.982, 13.6] | 0.0002 |
|  | Permethrin / Untreated | 2.83 [0.878, 9.14] | 0.0935 |

Supplementary Table 20: Among genotypes comparison of feeding duration for non-KD mosquitoes

| treatments | contrast | Hazard-Ratio [95% CI] | p-value |
| --- | --- | --- | --- |
| Permanet | RR / RS | 2.8105578 [0.7837099, 10.079284] | 0.13967169 |
|  | RR / SS | 3.8940624 [1.1198755, 13.540543] | 0.02849405 |
|  | RS / SS | 1.3855123 [0.4060183, 4.727975] | 0.80771517 |
| Olyset | RR / RS | 0.4674352 [0.138417, 1.578532] | 0.30808746 |
|  | RR / SS | 1.1864242 [0.3512593, 4.007303] | 0.9420286 |
|  | RS / SS | 2.5381578 [0.5708977, 11.284413] | 0.30878458 |
| Untreated Net | RR / RS | 2.1147017 [1.1120872, 4.021234] | 0.01735618 |
|  | RR / SS | 1.9475102 [1.0198706, 3.718899] | 0.04163654 |
|  | RS / SS | 0.9209385 [0.5102817, 1.662077] | 0.94278586 |

Supplementary Table 21: Among treatments comparison of weighted volume of bloodmeal for non-KD mosquitoes

| genotype | contrast | Difference [95% CI] | p-value |
| --- | --- | --- | --- |
| RR | Deltamethrin - Permethrin | 0.0248 [-0.91, 0.96] | 0.9978 |
|  | Deltamethrin - Untreated | -1.4802 [-2.36, -0.596] | 0.0004 |
|  | Permethrin - Untreated | -1.505 [-2.21, -0.805] | <.0001 |
| RS | Deltamethrin - Permethrin | 0.1193 [-1.12, 1.359] | 0.971 |
|  | Deltamethrin - Untreated | -2.0481 [-3.01, -1.085] | <.0001 |
|  | Permethrin - Untreated | -2.1675 [-3.23, -1.103] | <.0001 |
| SS | Deltamethrin - Permethrin | -0.1725 [-1.43, 1.082] | 0.9436 |
|  | Deltamethrin - Untreated | -1.0424 [-1.83, -0.255] | 0.0059 |
|  | Permethrin - Untreated | -0.8699 [-2.01, 0.271] | 0.1718 |

Supplementary Table 22: Among genotypes comparison of weighted volume of bloodmeal for non-KD mosquitoes

| treatment | contrast | Difference [95% CI] | p-value |
| --- | --- | --- | --- |
| Permanet | RR / RS | -0.419733 [-1.566527415, 0.7270613] | 0.65962081 |
|  | RR / SS | -0.6966876 [-1.742856362, 0.3494811] | 0.25650731 |
|  | RS / SS | -0.2769546 [-1.391801099, 0.8378919] | 0.82535704 |
| Olyset | RR / RS | -0.3252302 [-1.377845445, 0.727385] | 0.7338532 |
|  | RR / SS | -0.8939897 [-2.078087154, 0.2901078] | 0.17790592 |
|  | RS / SS | -0.5687595 [-1.967860526, 0.8303416] | 0.60058882 |
| Untreated | RR / RS | -0.9876998 [-1.728401344, -0.2469982] | 0.00642833 |
|  | RR / SS | -0.2588998 [-0.921484071, 0.4036844] | 0.61850042 |
|  | RS / SS | 0.7288 [0.001148454, 1.4564515] | 0.0495624 |
